# The environment topography alters the transition from single-cell populations to multicellular structures in *Myxococcus xanthus*

**DOI:** 10.1101/2021.01.27.428527

**Authors:** Karla C. Hernández Ramos, Edna Rodríguez-Sánchez, Juan Antonio Arias del Angel, Alejandro V. Arzola, Mariana Benítez, Ana E. Escalante, Alessio Franci, Giovanni Volpe, Natsuko Rivera-Yoshida

**Author notes:** Both authors equally contributed to this work (Karla C. Hernández Ramos, Edna Rodríguez-Sánchez).

## Abstract

The social soil-dwelling bacteria Myxococcus xanthus can form multicellular structures, known as fruiting bodies. Experiments in homogeneous environments have shown that this process is affected by the physico-chemical properties of the substrate, but they have largely neglected the role of complex topographies. We experimentally demonstrate that the topography alters single-cell motility and multicellular organization in M. xanthus. In topographies realized by randomly placing silica particles over agar plates, we observe that the cells’ interaction with particles drastically modifies the dynamics of cellular aggregation, leading to changes in the number, size and shape of the fruiting bodies, and even to arresting their formation in certain conditions. We further explore this type of cell-particle interaction in a minimal computational model. These results provide fundamental insights into how the environment topography influences the emergence of complex multicellular structures from single cells, which is a fundamental problem of biological, ecological and medical relevance.

## Introduction

Biological organisms exhibit complex collective behaviors heavily influenced by their intrinsic properties and their interaction with the environment. In the case of microorganisms, experiments have demonstrated that such behaviors are crucially affected by cellular properties such as shape, density, and motion dynamics (Starruß, et al., 2012; Be’er & Ariel, 2019; Nishiguchi, 2020). When considering the environment, the substrate’s physical properties have been shown to impact microorganisms survival, motility and collective behaviour (Velicer et al., 1998; Persat et al., 2015; Rivera-Yoshida et al., 2018; Bidossi, et al., 2020; Velic et al., 2020). However, those experimental efforts have largely neglected the complexity of the environment where the microorganisms live, favoring experimental conditions that greatly oversimplify ecologically meaningful contexts, partly due to the need to standardize experimental designs (Gilbert, 2001; Sultan, 2003; Robert, 2004; Rivera-Yoshida, et al., 2020). Recent work in the field of active matter has demonstrated that the topography of the environment can have a major influence on the motion and behavior of nonliving active particles (Volpe et al., 2011; Bechinger et al., 2016; Volpe & Volpe, 2017) and single bacterial cells (Frangipane, et al., 2019; Meel et al, 2012; Makarchuk et al., 2019; Velic et al., 2020), as well as on some collective bacterial phenomena (Be’er, et al. 2009; Persat et al., 2015; Lowery & Ursell, 2019; Rivera-Yoshida et al., 2018, 2019). Nevertheless, it is still an open question how complex topographies influence microorganism motility and behaviour throughout their levels of organization, from single cells to multicellular aggregates and structures.

*Myxococcus xanthus* is a motile rod-shaped soil-dwelling bacterium that glides across surfaces. It is a model system to study the transition to multicellularity, which is one of the most important transitions in the evolutionary history of life (Whitworth, 2008; Arias Del Angel et al., 2017, 2020). As part of its life cycle, *M. xanthus* exhibits different types of collective behaviour (Zusman et al., 2007; Zhang et al., 2012; Thutupalli et al., 2015; Muñoz-Dorado et al., 2016). In particular, when nutrients are depleted, *M. xanthus* transits into a developmental stage characterized by cellular aggregation, which culminates in the formation of densely packed multicellular structures called fruiting bodies, containing up to 10^6^ cells, where cell differentiation into spores takes place (Whitworth, 2008; Muñoz-Dorado et al., 2016). This developmental process involves different levels of organization, from single cells, to motile cell groups of different sizes, and finally to sedentary aggregates of millions of cells. At the onset of starvation, bacteria come together by colliding and following the slime trails left by other bacteria, thus forming motile aggregates that tend to increase in size and density (Kaiser 2003; Zusman et al., 2007; Whitworth, 2008; Thutupalli et al. 2015; Zhang et al. 2012). These aggregates can turn into streams, which sterically confine the cells and eventually permit the formation of three-dimensional stacks and fruiting bodies (Zhang et al., 2012; Thutupalli et al., 2015; Copenhagen et al., 2020; Figure 1).

**Figure 1.**
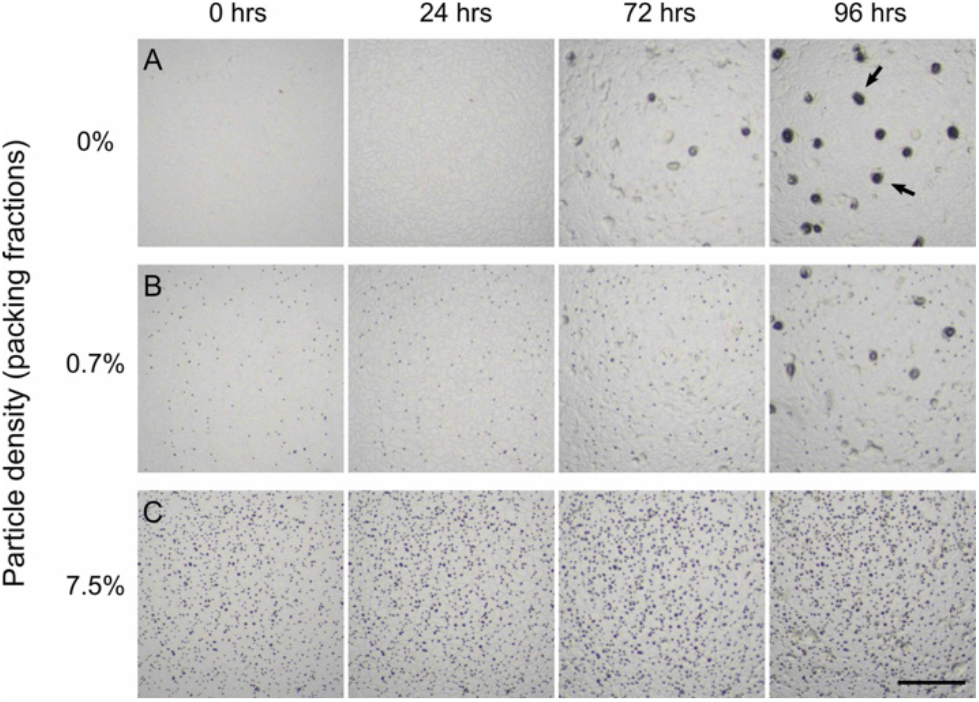
*M. xanthus* fruiting body development in homogeneous and heterogeneous topographies. *M. xanthus* cells come together to develop multicellular fruiting bodies, which can be identified as dark spots on agar plates. **a:** On a homogeneous substrate and at a cellular density of 0.01 OD, fruiting bodies start forming at 72 hrs and are completely mature by 96 hrs (some mature fruiting bodies are indicated by the arrows). **b**-**c:** Even a relatively small amount of silica particles randomly distributed over the agar surface can, **b**, hinder (0.7% particle packing fraction) or, **c**, completely prevent (7.5% particle packing fraction) the fruiting-body formation. The scale bar corresponds to 1 mm.

Various concepts from the field of soft and active matter have already been employed to understand the mechanisms underlying *M. xanthus* transition to multicellularity (Vicsek & Zafeiris, 2012; Starruß, et al., 2012; Bahar et al., 2014; Thutupalli et al., 2015; Liu et al., 2019, Arias Del Angel et al., 2020). For example, the dynamic formation, shrinkage and growth of cellular aggregates that ultimately produce fruiting bodies has been to some extent described as a droplet formation process in thin liquid films (Bahar et al., 2014) and as a phase separation driven by cells that change their motility over time (Thutupalli et al., 2015; Liu et al., 2019). Furthermore, the transition to the various phases that characterize *M. xanthus* development has been proposed to result from the regulation of motility factors, which are usually associated with genetic or strain-specific features of the cells, namely their speed, reversal frequency, and slime production rate (e.g. Morgan et al., 2010; Starruß, et al., 2012; Cotter et al., 2017). However, also some environmental properties can have a major influence on the collective behavior of *M. xanthus*. For example, the chemical and physical properties of the substrate (e.g., stiffness and tension) significantly affect the behaviour of *M. xanthus* and its multicellular development in terms of its fruiting body number, size, shape and spatial distribution (Stainer, 1942; Fontes & Kaiser, 1999; Meel et al., 2012; Rivera-Yoshida et al., 2018; Lemon et al., 2018). Moreover, although the natural populations of *M. xanthus* live in highly heterogeneous soil environments, the role of the substrate properties in *M. xanthus* motility and development has been studied only on smooth agar substrates. Thus, the question of how the environmental topography comes into play at different scales during unicellular-to-multicellular transitions, specifically in *M. xanthus*, remains to be explored.

Here, we experimentally demonstrate that the environment topography alters both single-cell motility and multicellular organization in *M. xanthus* colonies. Specifically, we demonstrate the effect of heterogeneous topographies, created by randomly placing silica particles over agar plates, on cellular motility, as well as on the number, size and shape of multicellular fruiting bodies. We find that the particles attract some individual cells, whose trails are then followed by other cells. This results in the sequestration of a sufficiently large number of cells to alter the formation of aggregates, effectively hindering the formation of multicellular fruiting bodies, specially at low cellular densities. We support our results with numerical simulations of early aggregation and discuss them in terms of the interplay between the physical and the biological processes involved in the development of multicellular fruiting bodies under ecologically meaningful conditions.

## RESULTS

### The substrate topography alters *M. xanthus* development and fruiting body formation

In the homogenous environment provided by a smooth agar plate, the *M. xanthus* colonies reach their complete development, fully maturing their fruiting bodies, in 96 hrs, as shown in Figure 1a. The fruiting bodies can be recognized by their shape and size, and by their dark color conferred by the presence of differentiated spores (see, e.g., arrows in Figure 1a). To determine the contribution of the environment topography to the collective dynamics of *M. xanthus*, we realized some complex environments by randomly distributing 10 μm diameter silica particles onto flat agar substrates. Specifically, we assayed the WT-DZF1 *M. xanthus* strain at six cellular densities (measured by their optical density (OD) at 550 nm:cellular densities: 0.01, 0.02, 0.06, 0.1, 0.3, 0.7 OD550) and seven different topographic conditions (measured by the particle area packing fraction, i.e., the fraction of the substrate area covered by the particles: 0%, 0.7%, 4.2%, 7.5%, 24%, 36%, 45%; see the Experimental growth and developmental conditions on heterogeneous substrates in Methods). In each condition, we took micrographs at 0, 24, 72 and 96 hrs (see the Macroscopic experimental measurement and data analysis section in Methods). Even a small amount of particles was sufficient to drastically modify *M. xanthus* multicellular development: a packing fraction of just 0.7% was sufficient to hinder the development of the fruiting bodies, as can be seen from their greatly reduced number in Figure 1b, while a larger packing fraction of 7.5% completely prevented their development, as can be seen from their absence in Figure 1c.

The development of fruiting bodies as a function of cellular density and particle density is shown in Figure 2a and in supplementary video 1. From this phase diagram, it is clear that the formation of fruiting bodies decreased both by decreasing the cellular density or by increasing the particle packing fraction. Strikingly, for low cellular densities and high particle packing fractions, a complete arrest of fruiting-body formation occured (lower left corner of Figure 2a, encircled by the pink line). If they managed to form, the number of fruiting bodies increased with the cellular density for each environmental topography (Figure 2b). Interestingly, the average size of the largest fruiting bodies decreased as the cellular density increased, while they featured a weaker dependency on the packing fraction of particles (Figure 2c). By contrast, the average size of the smallest fruiting bodies decreased as the particle packing fraction increased, while featuring a weaker increase as the cellular density increased (Figure 2d). Finally, we observed that the shape of the fruiting bodies became more elongated as the particle packing fraction increased (Figure 2e, the fruiting-body shape is measured as circularity *c =* 4π * *area/perimeter*^*2*^, where *c*=0 indicates an elongated shape and *c*=1 a perfect circle). These results demonstrate that the cellular density and the environment topography jointly affect fruiting-body maturation, number, size and shape, which are important developmental traits upon which natural selection may act.

**Figure 2.**
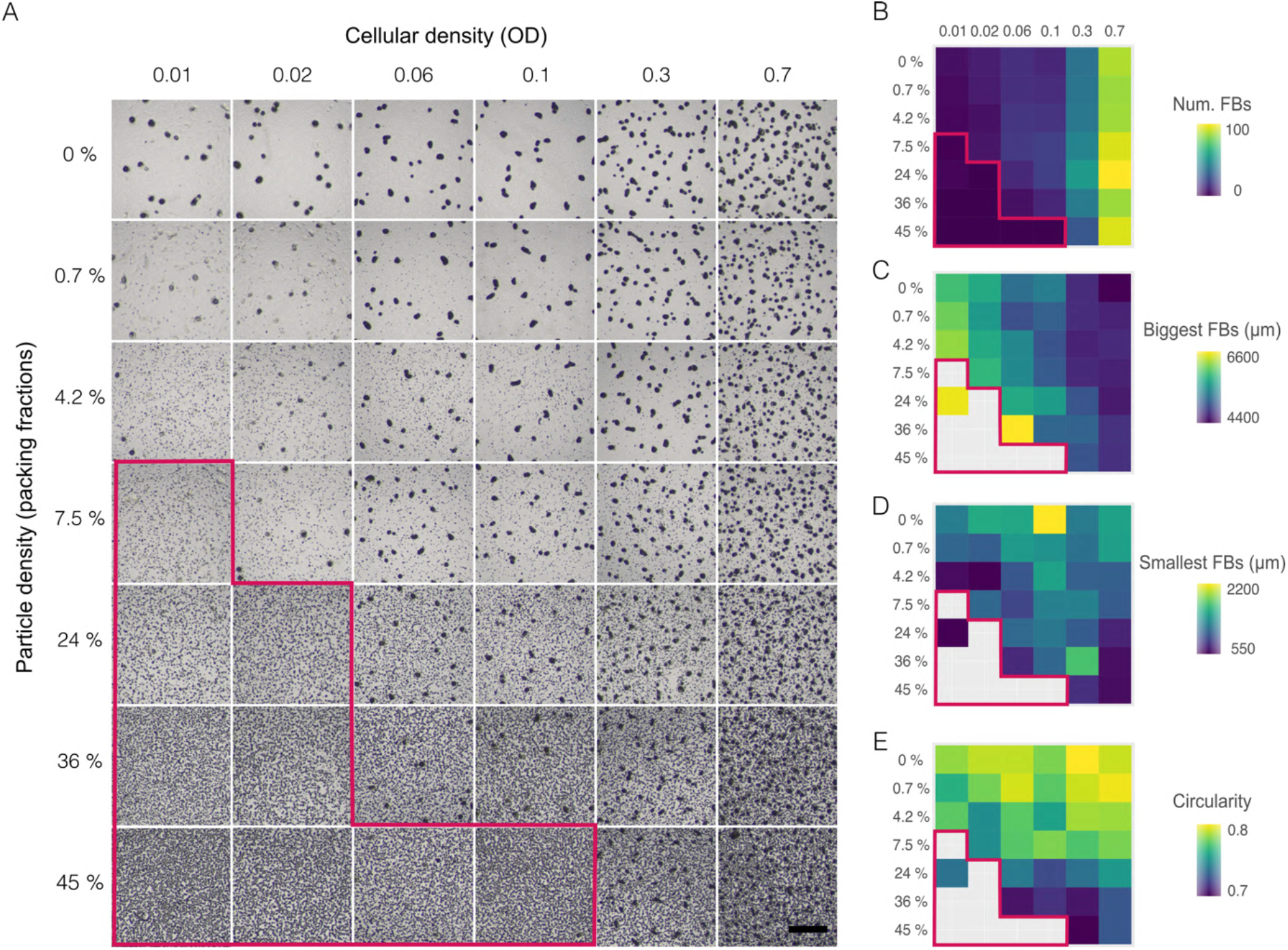
Fruiting-body formation dependence on cell and particle density. a: Micrographs at 96 hrs of *M. xanthus* populations generated over agar substrates at different particle densities (0%, 0.7%, 4.2%, 7.5%, 24%, 36%, 45% packing fractions) and at different cellular densities (0.01, 0.02, 0.06, 0.1, 0.3, 0.7 optical densities (OD) at 550 nm). For each condition, the micrograph shows the central section of the population. The large dark spots are the mature fruiting bodies (FBs), while the small dots in the background are the silica particles dispersed over the agar substrate (the scale bar corresponds to 1 mm). **b**-**e**: Mean variation of fruiting-body developmental traits as a function of cellular and particle density: **b** number, **c** average size of the five biggest fruiting bodies, **d** average size of the five smallest fruiting bodies, and **e** circularity as a shape descriptor. In all cases, the region within the pink line indicates the set of conditions in which fruiting-body formation is completely arrested. See also supplementary video 1.

### The particles disrupt the early aggregation dynamics by attracting and sequestering cells

To understand the microscopic mechanisms underlying the changes induced in the multicellular organization of *M. xanthus* by heterogeneous topographies, we studied the local cell-particle interaction during the early stages of aggregation. *M. xanthus* populations at intermediate cellular and particle densities (0.1 OD550, 24%, Figure 3a) were recorded for 2 hrs and then tracked in order to obtain the spatial coordinates and speed of the individual cells (Figures 3b,c; supplementary video 2; see the sections of Microscopic local observation and Tracking analysis of individual cells in the Methods). Over the course of hours, individual and aggregated cells moved towards nearby particles. When reaching the particles, the cells got arranged tangentially to the particles within the aqueous menisci that were formed around the particles (gray rings surrounding particles in Figure 3a, defined between *r* and *r*_*p*_ in figure 3b). These menisci are formed by capillary action at water-air-solid interfaces (Figure 3b). These menisci form also around cells (Figure 3c), even though in this case they are much smaller in comparison to those formed around particles (cfr. Figure 3b). Individual cells also migrated and collided into growing aggregates with a nematic arrangement, which, in turn, fused with other nearby aggregates or moved towards particles (Figure 3c, supplementary video 2).

**Figure 3.**
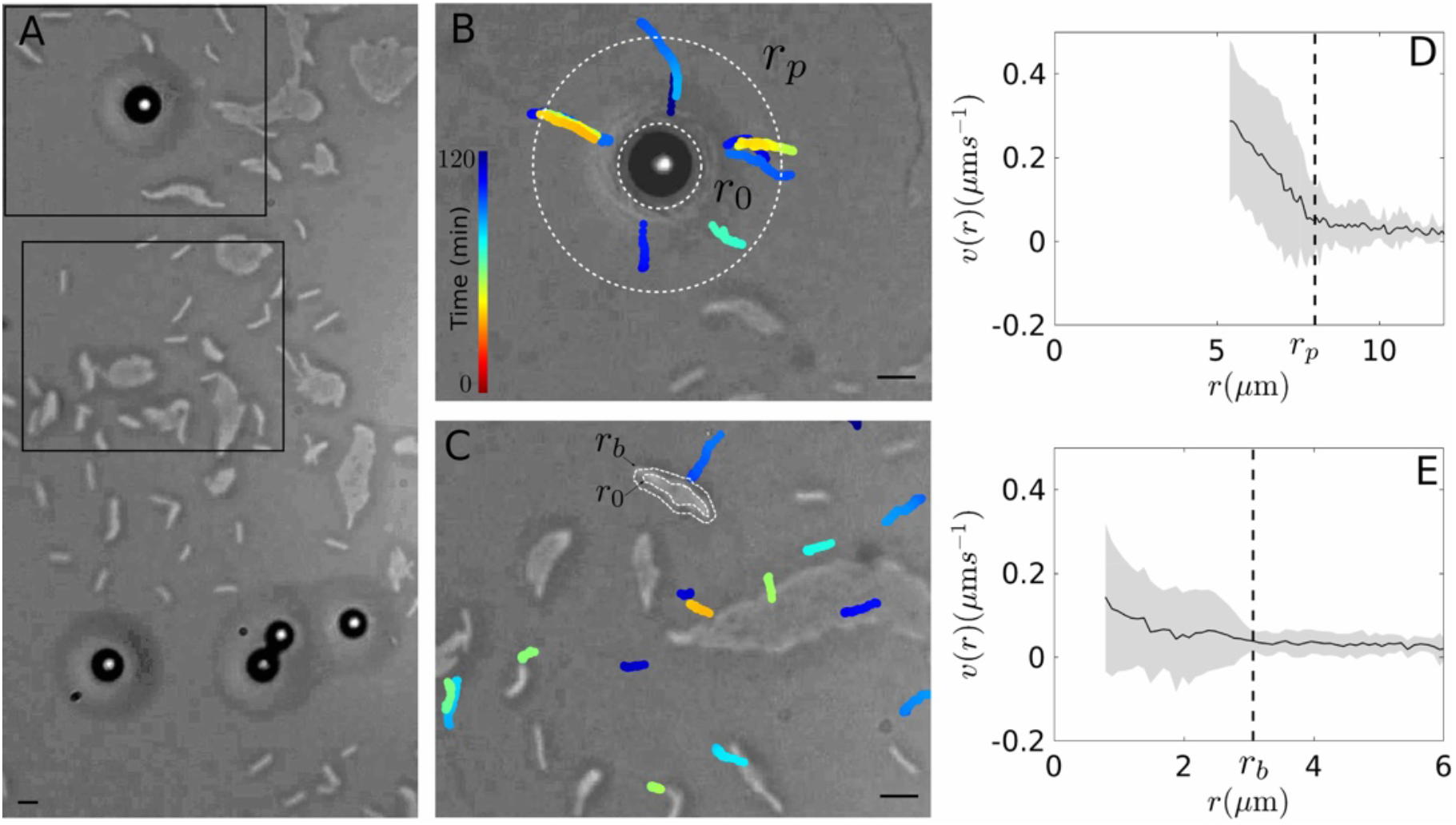
*M. xanthus* cell-cell and cell-particle local interactions during cellular aggregation. a: Initial configuration of some cells (elongated bright shapes) and particles (dark discs with bright spot). The gray rings around the silica particles are the aqueous menisci. **b:** Movement of cells near the particle during 2 hrs (corresponding to the top frame in **a**). The white dotted lines indicate the contact line of the aqueous meniscus (larger circle, *r*_*p*_) and the surface of the particle (smaller circle, *r0*). The coloured lines correspond to the trajectories of cells moving towards the particle; colors go from cold to warm according to the time when each position was recorded. Note that the cells’ trajectories tend to overlap, which reflects that the cells follow the slime trails left by other cells. **c:** Movement of cells far away from particles during 2 hrs (corresponding to the bottom frame in (a)). The white dotted lines indicate the contact line of the aqueous meniscus of the bacterial aggregate (larger outline, *r*_*b*_) and the surface of the aggregate (smaller outline, *r*_*0*_). The coloured lines correspond to the trajectories of cells moving towards cellular aggregates (note that some target aggregates moved from their initial position during the 2 hrs); the color code is the same as in **b. d-e:** Average speed of tracked cells while they approach, **d**, a particle and, **e**, other cells (corresponding to the trajectories in **b** and **c**, respectively). The gray-shaded areas represent one standard deviation. *r*_*0*_ and *r*_*p*_ corresponds to the surface and meniscus of the target particle or bacterial aggregate in **d** and **e**, respectively. The scale bars correspond to 10 μm. See supplementary video 2.

Cell migration could target either nearby particles or cellular aggregates (Figures 3b and 3c, respectively). Initially, cells featured an exploratory behavior characterized by short displacements in seemingly random directions, but as they came closer to a target, they started moving straight towards it (supplementary video 2). As quantified in Figures 3d,e, individual cells increased their speed exhibiting a drastic acceleration when they reached the triple-phase interface line of the particle’s or the cellular aggregates’ aqueous meniscus (*r*_*p*_ and *r*_*b*_ in Figure 3d,e). This strongly suggests that bacteria are pulled into the meniscus due to capillary attractive forces (Dalbe et al., 2011; Flury & Aramrak, 2017; Ho et al., 2019), which have been proposed to be a dominant factor of colloid deposition at interfaces and are one of the drivers of *M. xanthus* motility (Keller et al., 1983; Dworkin et al., 1983; Flury & Aramrak, 2017). Interestingly, the speed of cells moving towards particles (Figure 3d) was significantly larger than that of cells moving towards other cells (Figure 3e). Moreover, since the particle meniscus is larger than the cell meniscus, particles exerted a greater attraction over a larger area. We also observed that, once cells entered a particle’s meniscus, they remained there for the rest of the time (Figure 3b,c; supplementary video 2). Finally, some cells left a visible slime trail of their trajectory, which was later followed by other cells (Figure 3b,c; supplementary video 2). The existence of this trail-marking and trail-following mechanism in *M. xanthus* motility is well known (Kaiser 2003; Gloag et al., 2016; Muñoz-Dorado et al., 2016). Over time, the number of trails approaching a particle increased, which in turn further increased the effective strength and domain of the attraction exerted by the particle. Consequently, the space between particles was gradually depleted of cells, the majority of which accumulated around the particles (Figure 3b; supplementary video 2). This pronounced trapping effect does not occur with bacteria-bacteria interactions since individual and aggregated bacteria are always moving and reorganizing during this early stage, inhibiting the formation of persistent trails. We did not find evidence that the cells surrounding the particles, although somehow aggregated, could give rise to fruiting bodies.

These observations show that the heterogeneous substrates we designed disturb the aggregation and developmental dynamics of *M. xanthus* because the cells are attracted by particles with more strength than by other cells. Furthermore, as cells continuously modify the substrate by leaving trails that other cells can follow, these cellular trails reinforce the attraction of individual cells and groups of cells towards the particles, effectively enabling longer-distance attraction. Therefore, the combined action of attraction and subsequent trail-following renders large areas around the particles depleted of cells. As a consequence, the particles successfully compete with the cells, managing to attract a great number of cells and preventing the sequestered cells from forming fruiting bodies.

### Particle-cell attraction is crucial to alter cell aggregation

The experimental results presented in the previous sections show that *M. xanthus* cells are attracted to particles, mainly via capillary attraction forces and subsequent trail following, even more than they are attracted to other cells. To further test the overall effect of this cell-particle attraction on cell aggregation, we used a qualitative Glazier-Graner-Hogeweg computational model (Glazier et al., 2007), which is commonly employed to model cellular and developmental dynamics and has successfully reproduced different aspects of *M. xanthus* collective behaviour (Holmes et al. 2010; Swat et al., 2012; Balagam & Igoshin, 2015; Arias Del Angel et al., 2019). This model considers elongated and semi-flexible simulated cells capable of secreting trail-forming slime, and of adhering to other cells forming stable aggregates (see section on Computational Model in Methods; Figure 4a). The heterogeneous substrate was simulated adding particles at various densities (0%, 1%, 5%, 15%, 25% packing fraction; Figures 4b,c,d, respectively). The attractive effect of the aqueous meniscus observed in the experiment was modeled through a particle-cell attraction gradient given by a decreasing logarithmic function. The model parameters were selected on the basis of previous reports and of our own experimental observations (Table 1).

**Table 1.** Parameters used to simulate *M. xanthus* early aggregation. The referenced parameters are based on previously reported *M. xanthus* models or based on our experimental data.

| Parameter | Remark | Value | References |
| --- | --- | --- | --- |
| Lattice | lattice dimensions | 80 x 80 $\mu\text{m}$<br>160 x 160 pixels | This work |
| $T_m$ | amplitude of cell-membrane fluctuation | 10 | Swat et al. 2012 |
| $\lambda_L$ | cell elongation constriction value | 30 | This work |
| $L_\sigma$ | cell target length | 6 $\mu\text{m}$<br>12 pixels | Kaiser, 2003 |
| $C$ | connectivity penalty value | $10^{11}$ | This work |
| $\lambda_{cV}$ | cell volumen constriction value | 5 | Swat et al. 2012 |
| $V_{c\sigma}$ | cell target volume | 3 $\mu\text{m}^2$<br>12 pixels | Kaiser, 2003 |
| $\lambda_B$ | curvature constriction value | $10^6$ | Pelling et al. 2005 |
| $\lambda_S$ | slime following value | 10 | Swat et al. 2012 |
| $k_S$ | slime decay rate | 0.006 | Yu & Kaiser, 2007 |
| $F_S$ | uniform slime secretion | 0.9 | Yu & Kaiser, 2007 |
| $\lambda_P$ | cell-particle attraction value | 200 | This work |
| $k_P$ | particle attractive gradient,<br>particle domain of attraction | 0.003 | This work |
| $F_P$ | uniform renewal of particle<br>attractiveness | 0.9 | This work |
| $J_{cc}$<br>$J_{cp}$ | cell-cell adhesion<br>cell-particle adhesion | 10<br>7; 50 (attractive;<br>non-attractive<br>particles) | Swat et al. 2012<br>and this work |
| $J_{pp}$ | particle-particle adhesion | 50 | |
| $J_{pm}$ | particle-medium adhesion | 10 | |
| $J_{mm}$ | substrate-medium adhesion | 0 | |
| $J_{mc}$ | substrate-cell adhesion | 20 | |

**Figure 4.**
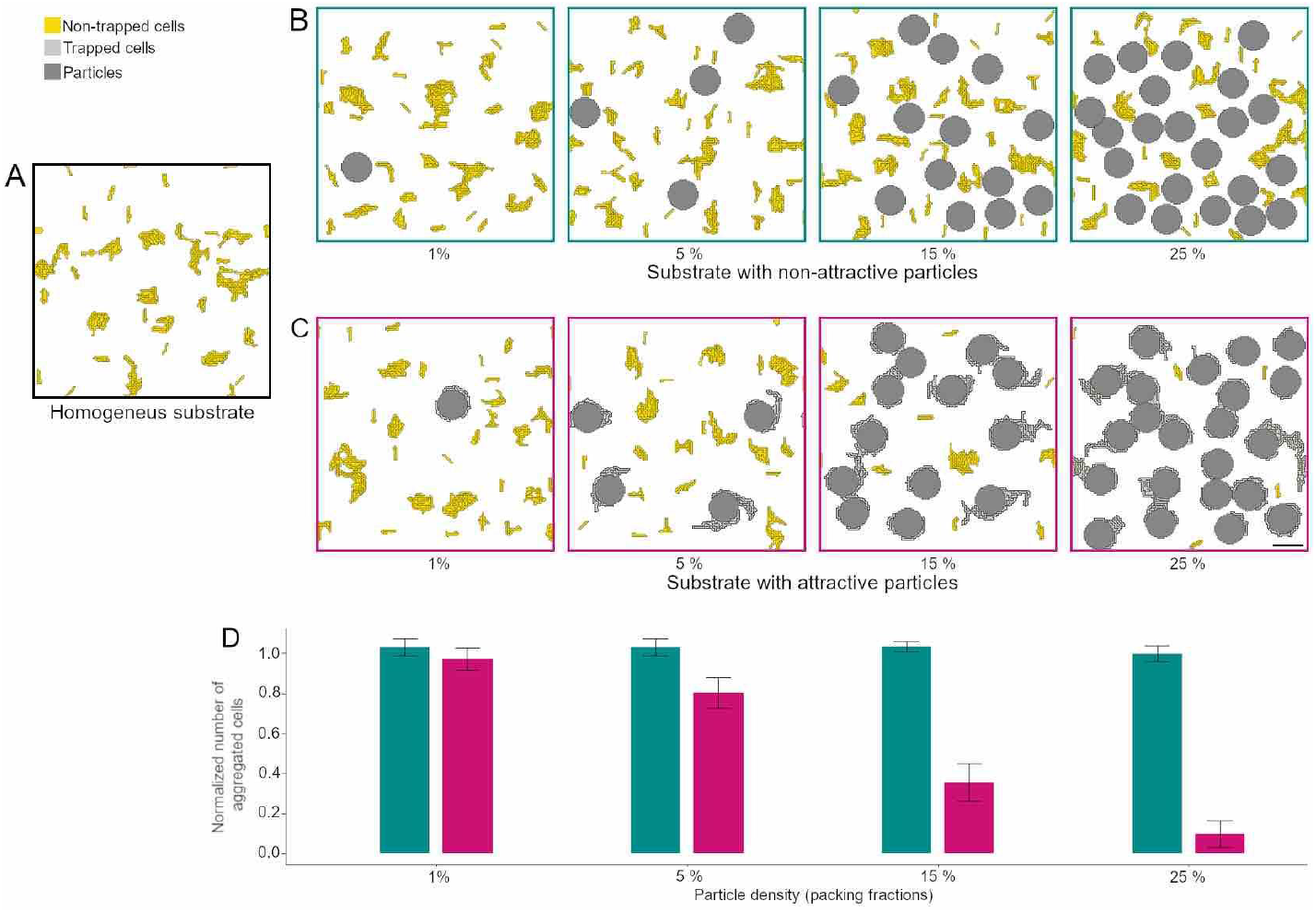
Effect of particle attraction in simulations of cellular aggregation. a: Simulation of *M. xanthus* cellular aggregation on a smooth and homogeneous substrate at 5000 Monte-Carlo steps. **b**-**c**: Corresponding simulation on a heterogeneous substrate varying the packing fraction (1%, 5%, 15%, 25%) of **b** non-attractive and **c** attractive particles. Particles are shown in gray, cells not trapped by particles in yellow, and cells trapped by particles in light-gray. The scale bar corresponds to 10 µm. **d:** Quantification of the number of aggregated cells, normalized to the aggregation numbers on the homogeneous substrate. The number of cells that are not trapped by particles drastically diminishes as the density of attractive particles increases, while it remains almost unchanged as the density of non-attractive particles increases. See supplementary video 3.

We simulated three types of substrate: i) a smooth substrate with no particles (Figure 4a), ii) a substrate with randomly distributed particles acting as steric obstacles (without attractive force) (Figure 4b), and iii) a substrate with randomly distributed particles attracting individual or aggregated cells (Figure 4c). In each one of these substrates, we introduced a population of simulated cells and allowed them to organize until steady aggregates were formed (around 5,000 Monte-Carlo steps). We performed ten replicates for each condition and counted the number of aggregated cells that were not trapped by the particles (Figure 4d). In the absence of particles (Figure 4a), simulated cells spontaneously aggregated in structures similar to those seen experimentally (Figure 3). We took the average number of aggregated cells in this smooth homogenous substrate as a reference for the scenarios with attractive and non-attractive particles. When non-attractive particles were added to the environment (Figures 4b,d, supplementary video 3), the average number of aggregated cells remained almost the same as in the reference case, with a slight decrease when the particle density increased to 25% of the area. In contrast, when attractive particles were added (Figures 4c,d, supplementary video 3), the average number of aggregated cells that were not trapped by a particle sharply decreased with particle density. While the features of heterogeneous media are often considered solely as obstacles for living active particles (e.g. Frangipane et al., 2019; Makarchuck et al., 2019; Rahmani et al., 2020), our simulation results support that attractive particles may sequester nearby cells, diminishing the local cellular density in the remaining area and hindering or completely preventing their fusion into bigger aggregates that could eventually give rise to fruiting bodies. More generally, attractive particles like those constituting the topography of our experimental setup establish a qualitatively different interaction with cells compared to steric obstacles, resulting in completely different dynamics and behaviours.

## DISCUSSION

We studied the development of the multicellular bacterium *M. xanthus* as a function of different cellular densities on substrates with varying environmental topographies, artificially built with randomly-deposited silica particles. We observed that the environment topography had a strong impact on the development of its multicellular fruiting bodies. We found that fewer fruiting bodies were formed as the particle packing fraction increased. This trend was particularly pronounced at low bacterial densities, for which the formation of fruiting bodies could even be completely inhibited, as shown in Figures 2a-b. Moreover, the presence of particles was associated with the formation of smaller fruiting bodies than in homogeneous substrates (Figures 2c-d), and with the change in fruiting-body shape from circular to elongated (Figure 2e). Our results are in line with previous works reporting that the substrate physical properties are able to modify the cellular spatial arrangement over the substrate (Stainer, 1942; Fontes & Kaiser, 1999; Veening et al., 2006; Meel et al., 2012; Lemon et al., 2018; Andac et al., 2019; Rivera-Yoshida et al., 2018, 2019), although these previous studies had not addressed the impact of the substrate topography on the emergence of multicellular structures. Considering that *M. xanthus* dwells in highly structured soil environments, it is of crucial importance to consider the fact that physical properties associated with heterogeneous topographies can affect the development of its multicellular structures, even arresting it in certain conditions.

When we looked into the detailed dynamics of cells at the onset of multicellular organization, we found that the particles used to generate the random topographies attracted the cells. Our direct observations of the cells’ trajectories moving toward particles or cellular aggregates showed that a drastic increase in their speed coincided with the reaching of the contact line of their target’s menisci (Figure 3d,e); this suggests that capillary forces mediate such attraction. This is in agreement with previous works proposing that capillary forces play a dominant role on colloid deposition and transport at solid-air-water interfaces (Bai et al., 2017; Flury & Aramrak, 2017; Bai et al., 2020), as well as with previous models and observations pointing to surface tension as a force driving *M. xanthus* motility (Dworkin, 1983; Dworkin et al., 1983; Keller et al., 1983). Indeed, these observations are in line with previous findings showing that *M. xanthus* gliding motility is associated with surface tension, which facilitates cellular adhesion to the substrate and generation of pushing forces (Keller et al., 1983; Dworkin et al., 1983; Nan & Zusman, 2011; Ducret et al., 2012), Importantly, the magnitude and range of the attraction exerted by the particles was larger than the attraction exerted by other cells (Figure 3; supplementary video 2), inhibiting the cells from forming large cell-only aggregates, which are a precursor of the fruiting bodies. While our results highlight the potential effect of capillary forces on bacterial motility and aggregation, these forces may have contrasting effects depending on the type of media and cells, and may interact in complex ways with other physical and biological processes (Flury & Aramrak, 2017). Further experimental and simulation efforts could help understand their role in different physical contexts, for example, employing different substrates, humidity conditions, and particle sizes or materials.

Our results reveal also a synergy between capillary attraction forces and the slime-trail following typical of *M. xanthus* motility (Kaiser 2003; Gloag et al., 2016; Muñoz-Dorado et al., 2016). Because cells leave behind slime trails as they are attracted toward a particle, an increasing number of trails pointing toward the particle meniscus build up over time. This self-reinforcing mechanism gradually increases both the net attraction strength and the attraction domain of particles over cells. In Nature, this mechanism could underlie different types of attractive interactions among living and non-living elements of the environment. From an ecological perspective, our observations highlight the role of living active matter in continuously shaping its environment. The resulting bidirectional interaction involves several physical and biological interplaying aspects.

We have now shown that fruiting-body formation can be arrested because the cells are sequestered by the particles. It is known from the literature that, as part of multicellular development, cells form stacks that eventually lead to the three-dimensional organization of fruiting bodies (Thutupalli, et al., 2015; Copenhagen et al., 2020). Furthermore, models for cellular differentiation within fruiting bodies suggest that spores are formed after the accumulation of certain molecules at the center of these cellular aggregates (Mora Van Cauwelaert, et al., 2015; Arias Del Angel et al., 2019). Taking this into account, we speculate that the cellular sequestration by particles prevents both cellular differentiation and formation of three-dimensional structures. This is also in agreement with the lack of evidence for fruiting-body formation around particles in our experimental setup, despite the local accumulation of cells.

The mechanism of cell sequestration that we postulate requires attraction exerted by the particles on the cells, in contrast with mechanisms involving particles that act just as steric barriers or obstacles (Frangipane et al., 2019; Rahmani et al., 2020). We have investigated this aspect by developing a computational model to compare the effect of particles with and without an attractive field on the motility and aggregation of simulated cells (Figure 4a, supplementary video 3). This model gives rise to qualitatively different patterns of aggregation for attractive and non-attractive particles, whose trends coincide with those observed in our experiments: while non-attractive particles act only as obstacles that increase the local cellular density (Figure 4b), attractive particles sequester cells and prevent the formation of aggregates that could eventually come together to form fruiting bodies (Figure 4c). These simulations highlight the important role of cell-particle attractive interactions in determining the trajectories and spatial arrangement of groups of cells, biofilms and other active materials.

In conclusion, we have shown that a heterogeneous topography greatly affects the formation of multicellular *M. xanthus* fruiting bodies, and have provided empirical and numerical evidence for an underlying mechanism based on the sequestration of cells. Specifically, we postulate a combination of physical and biological processes that lead to cell sequestration by particles and the concomitant alteration of multicellular development into fruiting bodies, especially at low cellular densities. This contributes to our understanding of how *M. xanthus* cells can respond to the physical attributes of their environment, and how the intricate organism-environment interactions are shaped. More generally, this work advances our understanding of the role of the environment topography not only on the general organization principles of active particles, but also on the origin and development of multicellular aggregates in complex ecological contexts.

## METHODS

### Experimental growth and developmental conditions on heterogeneous substrates

Following the protocol described in Yang & Higgs (Yang & Higgs, 2014), DZF1 strain was taken from a frozen stock by spotting 50 μl onto a casitone yeast extract (CYE) agar plate (1% Bacto Casitone, 10 mM Tris-HCl (pH 7.6), 0.5% yeast extract, 10 mM MOPS (pH 7.6) and 4 mM MgSO4) and incubated at 32°C for 2 days. Cells from the resulting colony were transferred to 25 ml of CYE liquid medium and incubated at 32°C, shaking at 250 r.p.m. overnight. The culture dilution was grown from 0.1 OD550 until it reached 0.7 OD550 (nutrient-rich liquid culture) taking a sample each 0.1 OD550. Dilutions from the 0.1 OD550 sample were made to obtain 0.01, 0.02 and 0.06 OD550 samples. Prior to the bacteria development assays, cells were harvested by spinning them at 8000 r.p.m. for 5 min. The resulting pellet was washed with a TRIS phosphate magnesium medium (TPM) solution (10 mM Tris-HCl (pH 7.6), 1 mM K2HPO4, 8 mM MgSO4) and resuspended in 1/10th of the original volume. Fifteen microlitres were spotted onto the heterogeneous substrate. After the spots dried, the plates were incubated at 32°C for 96 h. 0.01 OD550 and 0.02 OD550 samples were conducted five times, while 0.06 OD550-0.7 OD550 samples were conducted in triplicate for statistical support.

Heterogeneous substrates were fabricated randomly distributing 10 μm diameter silica particles onto the flat agar TPM substrate. To this end, TPM agar plates were first prepared by filling each with 30 ml TPM/agar media (1.5% agar concentration) and storing them overnight at 32 °C before use. Then, ∼7.9 mg of dry silica particles were resuspended in 1 ml of deionized water. Dilutions of 2:3, 1:3, 1:10, 1:20 and 1:100 were obtained from this concentrated particle stock and 40 μl of each were spotted onto the TPM agar plates.

### Macroscopic experimental measurement and data analysis

To study the aggregation at the population level, micrographs of the resulting fruiting bodies were taken at 370.8 pixels/mm using a Leica m50 stereomicroscope with an Achro 0.63 objective lens and a Canon-EOS Rebel T3i camera. To avoid border effects, the fruiting bodies developed at the edge of the population were not considered. For image processing, micrographs were binarized into black/white images and phenotypic traits were measured using FIJI (ImageJ) software v. 2.0.0 (Schindelin et al., 2012). The changes in development due to the substrate heterogeneity and the cellular density were quantified and plotted by heatmaps for each trait (Figure 2b, c, d, e).

### Microscopic local observation

To examine the cell-cell and the cell-particle local interactions a sample at 0.1 OD550 cellular density and 24% particle density was observed using an optic microscope with an Olympus UPlanSApo 20x lens and recorded with a Basler acA 1600-20uc camera for 16 hr at 23 °C. To prepare the sample, 40 μl TPM/agar media (1.5% agar concentration) were deposited over a coverslip and, 10 min later, 3 μl of the cellular culture were spotted onto the TPM substrate. Ten minutes later, the sample was sealed using a coverslip and vacuum grease and it was then observed for 2 hrs at 23 °C at a sampling rate of 0.5 frames s^-1^.

### Tracking analysis of individual cells

For the tracking analysis shown in Figure 3, a semi-automatic tracking was performed following morphological properties of the cells with a homemade routine in Matlab. For each individual trajectory, the distance, speed and spatial coordinates were obtained.

### Computational model

To explore *M. xanthus* early aggregation in different topographies, we built a Glazier-Graner-Hogeweg model (also known as cellular Potts model) employing the CompuCell3D 3.7.8 platform (Glazier et al., 2007; Swat et al., 2012). Our model is based on *M. xanthus* experimental data and previously published models for this organism (Bahar et al., 2014; Holmes et al., 2010; Kaiser, 2003; Pelling et al., 2005; Starruß et al., 2007; Yu & Kaiser, 2007). The cellular properties and their interactions were implemented by a Halmitonian energy equation whose minimization, through the Monte-Carlo algorithm, drives the dynamics of the system (Table 1; Swat et al., 2012):

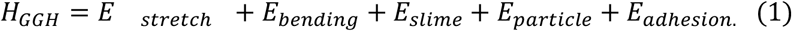

The *M. xanthus* rod-shape was represented by 12 continuous segments (Holmes et al., 2010) that hold their shape through the stretch energy

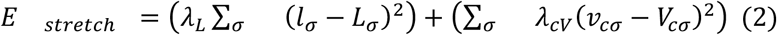

where the first term refers to the elongation *L* and the second to the volume *V* (area in our model). *l*_*σ*_is the current length of a cell *σ, L*_*σ*_ the reference length, and *λ* _*L*_ the elongation constriction value. The same notation applies to the second term of the equation. Cells breakdown was avoided by introducing a connectivity penalty value *C* (Merks et al., 2006). The simulated cells were randomly added on the substrate.

The cell semi-flexible movement was described by the bending energy avoiding abnormal contractions by

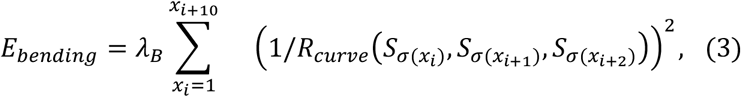

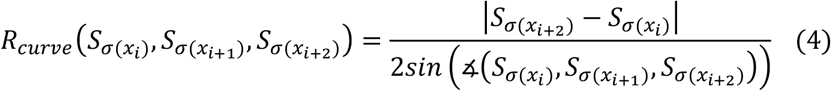

where *λ*_*B*_ is the curvature constriction value, and *R*_*curve*_ is the curvature radius of three consecutive cell segments *x*_*i*_, *x*_*i*+1_, *x*_*i*+2_ of a cell σ (Starruß et al., 2007).

The secretion of slime trails and their tracking by other cells was implemented using

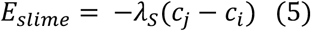

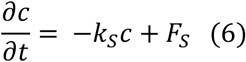

where *c*_*i*_ and *c*_*j*_ are the slime concentration at sites *i* and *j*, λ_*S*_ is the slime attraction coefficient, *F*_*S*_ is the uniform slime secretion and *k*_*s*_ is the logarithmic decay rate. Thus, the cells tended to move from lower to higher slime concentration sites.

To simulate the heterogeneous media, attractive particles were modeled through

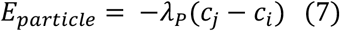

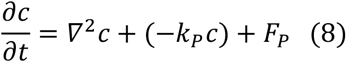

where *c*_*i*_ and *c*_*j*_ are the attraction values at sites *i* and *j*, λ_*P*_ is the particle attraction coefficient, *F*_*p*_ is the uniform renewal of the particle attractiveness and *k*_*p*_ is the attractive gradient of particles. 10 µm diameter particles were randomly added; the volume and position of the particles were static throughout the simulation (“freeze” function in CompuCell3D).

The adhesion of the model elements (cell, particle and substrate) was defined by

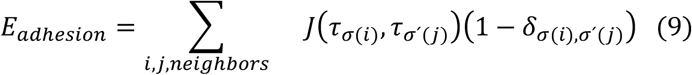

*J*represents the adhesion value between two contiguous pixels*i, j* that belong to a particular item σ,σ’, and an element τ. The second term prevents considering pixels that are not at the boundary of each item with a dirac function *δ*(Swat et al., 2012).

### Simulation execution and analysis

The model was executed for a cellular density of 0.03 cells/µm^2^ and 5 particle densities (0%, 1%, 5%, 15%, 25% packing fraction) simulating two scenarios, with and without cell-particle attraction. In each case, the total number of aggregated cells within and outside the particle attraction domain was obtained for all conditions. This was done for 10 replicas at 5,000 Monte-Carlo steps (Figure 4). We considered as biologically relevant aggregates those groups of cells formed by at least three cells. This parameter was chosen examining the frequency of the number of adhered cells at the beginning and at the end of the simulation without particles. Aggregates of three cells were infrequent at the beginning of the simulation and increased their frequency at the end, suggesting that their presence is not due to chance but to the aggregation process. Because we hypothesized that experimental aggregates within the particle menisci did not contribute to eventual fruiting bodies, we excluded those simulated aggregates. The number of aggregated cells was normalized with respect to the number of aggregated cells in homogeneous condition (Figure 4a).

Analyses were conducted in R (v. 4.0.2) using RStudio (Team R, 2014; RStudio Team, 2015). The ggplot2 package v. 3.0.0 was employed for visualization (Wickham, 2016).

## DATA AVAILABILITY

The dataset analyzed and the code generated for computational simulations during the current study are available in the GitHub repository, respectively:

https://github.com/laparcela/Myxo_heterogeneous_data_exp.git

https://github.com/laparcela/Myxo_heterogeneous_media_model/tree/master/Myxoliquid

## Acknowledgements

We thank Teresa Caudillo Estrada, Morena Avitia and Marco Tulio Solano De la Cruz for their technical assistance in the laboratory; James Glazier, Julio Belmonte and the CC3D team for technical support, as well as members of Laboratorio de Micromanipulación Óptica and LaParcela for insightful discussions. Financial support was provided by PAPIIT-DGAPA-UNAM IN102420 and IN111919, Conacyt Ciencia Básica A1-S-10610. N.R-Y. thanks POR EL APOYO DEL PROGRAMA DE BECAS POSDOCTORALES UNAM-DGAPA for a postdoctoral fellowship.

## Author contributions

K.C.H.R., N.R-Y. and A.V.A. performed the experiments and analysed the data; E.R.S. performed the simulations, with J.A.A.A’s help. K.C.H.R., N.R-Y., A.V.A., E.R.S., A.F., J.A.A.A., G. V. and M.B. discussed the experimental and simulations design and the data. N.R.Y., M.B. and G. V. wrote the article. All authors substantially revised the work and the final version of the article, except for J.A.A.A who passed away during the preparation of this paper.

