## Supplementary material for "The environment topography alters the transition from single-cell populations to multicellular structures in *Myxococcus xanthus*"

^3^ Current address: Posgrado en Ciencias Biológicas, Instituto de Geología, Universidad Nacional Autónoma de México, Mexico City, 04510, Mexico.

**SUPPLEMENTAL INFORMATION**

**Supplementary video 1. *M. xanthus* fruiting-body formation in homogeneous and heterogeneous topographies with varying cell density across time.**

Micrographs of *M. xanthus* populations generated over agar substrates at different particle densities (0%, 0.7%, 4.2%, 7.5%, 24%, 36%, 45% packing fractions; rows) and at different cellular densities (0.01, 0.02, 0.06, 0.1, 0.3, 0.7 optical densities (OD) at 550 nm; columns). For each condition, micrographs show the central section of the population at 0, 24, 72 and 96 hrs. The large dark spots are the mature fruiting bodies, while the small dots in the background are the silica particles dispersed over the agar substrate. Fruiting-body formation occurs earlier at low-to-mid-particle-density topographies (0-7.5%), compared to high-particle-densities topographies (24-45%) where development is inhibited at low cellular densities. The scale bar corresponds to 1 mm.

**Supplementary video2. *M. xanthus* cell-cell and cell-particle local interaction dynamics during cellular aggregation.** The time lapse shows the interaction between cells (elongated bright shapes) and particles (dark discs with bright spot) during 2 hrs. The gray rings around the silica particles and the gray shade surrounding the cellular aggregates are their respective aqueous menisci. Cells increase their speed when they reach the aqueous meniscus. The coloured lines correspond to the trajectories of cells moving towards particles or other cells; colors go from cold to warm according to the moment in which each position was recorded. The overlapping of some cells’ trajectories reflects that the cells follow the trails left by other cells. The scale bars correspond to 10 μm.

**Supplementary video 3. Early aggregation of *M. xanthus* simulated cells.** In **a** homogeneous topography and in **b-c** heterogeneous topographies at different packing densities of 1%, 5%, 15%, and 25% (from left to right) formed by **b** non-attractive and **c** attractive particles. Cells are shown in yellow and particles in gray. The cell aggregation is similar in the homogeneous environment and in the presence of non-attractive particles, even at the highest packing densities of the latter. Instead, in the presence of attractive particles, the formation of large aggregates decreases as the packing density increases. The sequence corresponds to simulations of 5000 Monte Carlo steps.
